# Deletion of Stim1 in hypothalamic arcuate nucleus Kiss1 neurons potentiates synchronous GCaMP activity and protects against diet-induced obesity

**DOI:** 10.1101/2020.09.09.289017

**Authors:** Jian Qiu, Todd L. Stincic, Martha A. Bosch, Ashley M. Connors, Stefanie Kaech Petrie, Oline K. Rønnekleiv, Martin J. Kelly

## Abstract

Kisspeptin (Kiss1) neurons are essential for reproduction, but their role in the control of energy balance and other homeostatic functions remains unclear. High frequency firing of hypothalamic arcuate Kiss1 (Kiss1^ARH^) neurons releases kisspeptin into the median eminence, and neurokinin B (NKB) and dynorphin onto neighboring Kiss1^ARH^ neurons to generate a slow excitatory postsynaptic potential (EPSP) mediated by TRPC5 channels that entrains intermittent, synchronous firing of Kiss1^ARH^ neurons. High frequency optogenetic stimulation of Kiss1^ARH^ neurons releases glutamate to excite the anorexigenic proopiomelanocortin (POMC) neurons and inhibit the orexigenic neuropeptide Y/agouti-related peptide (AgRP) neurons via metabotropic glutamate receptors. At the molecular level, the endoplasmic reticulum calcium-sensing protein stromal interaction molecule 1 (STIM1) is critically involved in the regulation of neuronal Ca^2+^ signaling and neuronal excitability through its interaction with plasma membrane calcium (*e*.*g*., TRPC) channels. 17β-estradiol (E2) downregulates *Stim1* mRNA expression in female arcuate neurons. Therefore, we hypothesized that deletion of *Stim1* in Kiss1^ARH^ neurons would increase neuronal excitability and their synchronous firing, which ultimately would affect energy homeostasis. Using optogenetics in combination with whole-cell recording and GCaMP6 imaging in slices, we discovered that the deletion of *Stim1* in Kiss1 neurons significantly increased the amplitude of the slow EPSP and augmented synchronous [Ca^2+^]i oscillations in Kiss1^ARH^ neurons. Deletion of *Stim1* in Kiss1^ARH^ neurons amplified the actions of NKB and protected ovariectomized female mice from developing obesity and glucose intolerance with high-fat dieting. Therefore, STIM1 appears to play a critical role in regulating synchronous firing of Kiss1^ARH^ neurons, which ultimately affects energy homeostasis.

**Significance Statement:** Hypothalamic arcuate kisspeptin (Kiss1^ARH^) neurons are essential for stimulating the pulsatile release of gonadotropin releasing hormone (GnRH) and maintaining fertility. However, Kiss1^ARH^ neurons appear to be a key player in coordinating energy balance with reproduction. The regulation of calcium channels and hence calcium signaling is critically dependent on the endoplasmic reticulum calcium-sensing protein stromal interaction molecule 1 (STIM1), which interacts with the plasma membrane calcium channels. We have conditionally deleted *Stim1* in Kiss1^ARH^ neurons and found that it significantly increased the excitability of Kiss1^ARH^ neurons and protected ovariectomized female mice from developing obesity and glucose intolerance with high-fat dieting.

## Introduction

Nutrition and reproduction are inextricably linked across all mammalian species, *i*.*e*., high circulating concentrations of 17β-estradiol (E2) during the late follicular phase of the reproductive cycle correlate with reduced food intake (Czaja, 1978; Asarian and Geary, 2006; Roepke et al., 2010). However, we are just beginning to understand the central mechanisms by which E2 feedback coordinates reproduction and energy balance (Castellano and Tena-Sempere, 2013; Nestor et al., 2014; Navarro, 2020). Kisspeptin neurons in the hypothalamic arcuate nucleus (Kiss1^ARH^ neurons) appear to be critical for coordinating these two homeostatic processes. Firstly, Kiss1 and its G protein-coupled receptor (GPR54) are essential for pubertal development and reproductive function (Kuohung and Kaiser, 2006). Mutations in Kiss1 or GPR54 cause hypogonadotropic hypogonadism in humans (De Roux et al., 2003; Seminara et al., 2003; Topaloglu et al., 2012), and deletion of Kiss1 or GPR54 causes defective sexual development and reproductive failure in mice (Seminara et al., 2003; d’Anglemont de Tassigny et al., 2007). These effects on fertility are directly dependent on Kiss1/GPR54 signaling in gonadotropin-releasing hormone (GnRH) neurons (Han et al., 2005; Pielecka-Fortuna et al., 2008; Zhang et al., 2008). Moreover, Kiss1 signaling appears to be also important for normal metabolism and glucose homeostasis. GPR54 deletion in female, but not male, mice causes severe obesity, reduced metabolism, glucose intolerance and hyperleptinemia (Tolson et al., 2014; Tolson et al., 2019). Also, Kiss1^ARH^ neurons are directly depolarized/excited by leptin (Qiu et al., 2011) and insulin (Qiu et al., 2014), so they are quite possibly the key neurons involved in conveying metabolic information to GnRH neurons.

High frequency optogenetic stimulation of Kiss1^ARH^ neurons expressing channel rhodopsin (ChR2) generates pulsatile release of LH (Clarkson et al., 2017). Kiss1^ARH^ neurons co-express neurokinin B (NKB) and dynorphin (Goodman et al., 2007; Navarro et al., 2009) and high-frequency firing (10-20 Hz) of these neurons co-releases NKB and dynorphin to coordinate the synchronous firing of the whole population of Kiss1^ARH^ neurons (Qiu et al., 2016). NKB binds to tachykinin 3 receptor (TacR3) in neighboring Kiss1^ARH^ neurons to activate canonical transient receptor potential 5 (TRPC5) channels to cause a robust depolarization (slow EPSP), whereas co-released dynorphin feeds back to bind to presynaptic κ-opioid receptors to limit the release of NKB to discrete bursts of activity (Qiu et al., 2016). The co-release of the two peptide neurotransmitters coordinates the synchronous firing of Kiss1^ARH^ neurons that drives the pulsatile release of GnRH into the median eminence (Qiu et al., 2016; Clarkson et al., 2017).

The activity of TRPC channels is modulated by stromal-interaction molecule 1 (STIM1), which is localized to the endoplasmic reticulum (ER) membrane of cells, and its N-terminal domain contains an EF-hand that senses changes in ER calcium concentrations and maintains intracellular Ca^2+^ homeostasis through store-operated Ca^2+^ entry (SOCE) (Salido et al., 2011). Upon depletion of ER Ca^2+^, STIM1 oligomerizes and then interacts with plasma membrane calcium (TRPC) channels (Yuan et al., 2007; Salido et al., 2011). Phosphorylation of STIM1 is required for oligomerization, and E2 inhibits the phosphorylation of STIM1 and its interaction with plasma membrane Orai and TRPC channels and hence store-operated Ca^2+^ entry (Yuan et al., 2007; Salido et al., 2011). Under normal physiological conditions, TRPC5 channels are coupled to plasma membrane receptors (Qiu et al., 2010; Qiu et al., 2014; Gao et al., 2017), but in cellular stressed states (*e*.*g*., inflammation, obesity) TRPC5 channels may associate with STIM1 to replete ER Ca^2+^ stores (Birnbaumer, 2009; Qiu et al., 2018b). E2 maintains the excitatory effects of insulin in POMC neurons, mediated by TRPC5 channel opening, by downregulating *Stim1* expression, thereby protecting against insulin resistance in obese females (Qiu et al., 2018b). E2 also downregulates *Stim1* expression in the ARH of female guinea pigs, indicating that this interaction is more widespread in the ARH. Therefore, we hypothesized that deletion of *Stim1* in Kiss1^ARH^ neurons would augment TacR3 mediated depolarization via TRPC5 channels to ultimately drive synchronous firing of the “pulse generator Kiss1^ARH^ neurons.

## Materials and Methods

### Animals

All animal procedures were conducted at Oregon Health and Science University (OHSU) according to the National Institutes of Health Guide for the Care and Use of Laboratory Animals and with approval from the OHSU Animal Care and Use Committee.

We used female mice in all of the experiments. *Kiss1*^*Cre:GFP*^ (v2) mice (Dr. Richard D. Palmiter; University of Washington; PMID: 29336844) (Padilla et al., 2018) were housed under constant temperature (21–23°C) and 12-h light, 12-h dark cycle schedule (lights on at 0600 and lights off at 1800 h), with free access to food (Lab Diets 5L0D) and water. *Kiss1*^*Cre:GFP*^ mice were used for viral injection to express ChR2 or GCaMP6s in Kiss1^ARH^ neurons or they were crossed with heterozygous *Ai32* mice (RRID:IMSR_JAX:024109, C57BL/6 background) purchased from The Jackson Laboratory. These *Ai32* mice carry the ChR2 (H134R)–EYFP gene in their Gt(ROSA)26Sor locus (Madisen et al., 2012). The gene is separated from its CAG promoter by a loxP-flanked transcriptional STOP cassette, allowing its expression in a Cre-dependent manner. To test for this we dispersed and harvested EYFP neurons in the ARH from *Kiss1*^*Cre:GFP*^*::Ai32* females and used single cell RT-PCR to determine *Kiss1* mRNA expression as described below and according to previous published methods (Bosch et al., 2013). Data from 126 ARH^EYFP^ neurons from 6 *Kiss1*^*Cre:GFP*^*::Ai32* females documented that 99% of the EYFP neurons expressed *Kiss1*.

To generate mice with conditional knockout of *Stim1* in Kiss1 neurons (*Stim1*^*kko*^), we first crossed Kiss1^Cre/+^ (v2) males (Padilla et al., 2018) with Stim1^loxP/loxP^ females (Jackson Laboratory Stock #023350, RRID:IMSR_JAX:023350, (Oh-hora et al., 2008)). This cross knocks out *Stim1* through excising exon 2 (Oh-hora et al., 2008) of the floxed *Stim1* gene in cells in which Cre is expressed under the control of a promoter specific for the expression of *Kiss1* (Padilla et al., 2018; Qiu et al., 2018a). The F1 mice produced were Kiss1^Cre/+^::Stim1^+/lox^ and Stim1^+/lox^. The F2 mice were generated by crossing these Kiss1^Cre/+^::Stim1^+/lox^ males with Stim1^loxP/lox^ females. Approximately 25% of the offspring were Kiss1^Cre/+^::Stim1^lox/lox^ such that Stim1 was deleted in Kiss1 cells (Stim1^KKO^), and all the Stim1 knock-out mice were seen at the expected frequency and viable throughout adulthood. We used Kiss1^Cre/+^ mice as controls. To increase the yield of Stim1 knock-out mice, we crossed Kiss1^Cre/+^::Stim1^lox/lox^ males with Stim1 ^lox/lox^ females. We maintained not only this strain but also the Kiss1^Cre/+^ strain at the same time. Genotypes for *Stim1* were determined using forward primer JAX#18885 (5’-CGA TGG TCT CAC GGT CTC TA-3’) and reverse primer JAX#18886 (5’-GCT CTG CTG ACC TGG AAC TA-3’), which distinguished between lox/lox, lox/+, and +/+ genotypes. Cre genotypes were determined using forward primer 5’-GCG GTC TGG CAG TAA AAA CTA TC3’- and reverse primer 5’-TTC CAT GAG TGA ACG AAC CTG G-3’, which distinguished between carriers and non-carriers of the Cre allele.

### Puberty onset and estrous cyclicity

To determine whether deleting *Stim1* in Kiss1-expressing neurons might impact fertility, we evaluated female *Stim1*^*kko*^ mice and wild type (WT) female littermates for pubertal onset and estrous cyclicity. For breeding, male and female mice were mated at 1:1, and the number of pups per litter was counted. The *Stim1*^*kko*^ mice showed similar fecundity as control mice. Puberty onset in females was assessed by monitoring for vaginal opening daily between 0900 and 1000 hr starting at 3 weeks of age. For estrous cycle studies, *Stim1*^*kko*^ and *Kiss1*^*Cre:GFP*^ female mice were group housed and were habituated to handling for at least one week by the same investigator prior to estrous cycle monitoring. Vaginal lavage was performed daily for 13 consecutive days between 0900 and 1000 hr. Cytology was evaluated using a light microscope and scored as diestrus, proestrus or estrus as previously described (Qiu et al., 2018a). The Number of estrous and diestrous days were counted for each animal and used for statistical analysis (Mann-Whitney U-test).

### Gonadectomy

At least 7 days prior to each experiment, ovaries were removed as described previously while under inhalant isofluorane anesthesia (Piramal Enterprises Limited, Andhra Pradesh, India) (Qiu et al., 2018a). Each mouse received analgesia (Carprofen; 5mg/kg; subcutaneous) immediately after a surgery for relief of postoperative pain.

### Metabolic Studies

For the metabolic studies, *Stim1*^*kko*^ and *Kiss1* littermate control females were ovariectomized at 2-4 months of age and put on a high fat diet (HFD; 45% kcal from fat; Research Diets, New Brunswick, NJ; D12451) for eight weeks. Mice were group housed (because of COVID-19 restrictions) and individually weighed every week. The evening prior to the glucose tolerance test (GTT), all mice were assessed for body composition (fat and lean mass) using an EchoMRI 4-in-1-500 Body Composition Analyzer (Houston, TX).

For GTT, age matched *Kiss1*^*Cre*^ and *Stim1*^*kko*^ mice were fasted overnight for 15-h, and baseline glucose levels measured with the aid of an Accu-Check Advantage blood glucose meter (Roche) using blood collected from the tail vein. All mice were then injected intraperitoneally with glucose (1 mg/g lean mass as determined by EchoMRI) in sterile PBS and blood glucose levels were measured 15, 30, 60, 90, and 120 min after injection. The glucose clearance (area under the curve) was calculated based on the glucose baseline levels at 0 min (Ayala et al., 2010).

### AAV delivery to Kiss1^Cre:GFP^ and Stim1^kko^ mice

Fourteen to twenty-one days prior to each experiment, *Kiss1*^*Cre:GFP*^ mice or *Stim1kko* mice (>60 days old) received bilateral ARH injections of a Cre-dependent adeno-associated viral (AAV; serotype 1) vector encoding ChR2-mCherry (AAV1-Ef1a-DIO-ChR2: mCherry) or ChR2-YFP (AAV1-Ef1a-DIO-ChR2:YFP, Dr. Stephanie L. Padilla; University of Washington; PMID: 25429312) or GCaMP6s (AAV9-Syn-Flex-GCaMP6s-WPRE-SV40; Addgene, # 100845-AAV9). Using aseptic techniques, anesthetized female mice (1.5% isoflurane/O_2_) received a medial skin incision to expose the surface of the skull. The glass pipette (Drummond Scientific #3-000-203-G/X; Broomall, PA) with a beveled tip (diameter = 45 μm) was filled with mineral oil, loaded with an aliquot of AAV using a Nanoject II (Drummond Scientific). ARH injection coordinates were anteroposterior (AP): −1.20 mm, mediolateral (ML): ± 0.30 mm, dorsoventral (DL): −5.80 mm (surface of brain z = 0.0 mm); 500 nl of the AAV (2.0 × 10^12^ particles/ml) was injected (100 nl/min) into each position, left in place for 10 min post-injection, then the pipette was slowly removed from the brain. The skin incision was closed using skin adhesive, and each mouse received analgesia (Carprofen; 5 mg/kg) for two days post-operation.

### Electrophysiology

Coronal brain slices (250 μm) containing the ARH from gonadectomized females were prepared as previously described (Qiu et al., 2003). Whole-cell, patch recordings were performed in voltage clamp and current clamp using an Olympus BX51W1 upright microscope equipped with video-enhanced, infrared-differential interference contrast (IR-DIC) and an Exfo X-Cite 120 Series fluorescence light source. Electrodes were fabricated from borosilicate glass (1.5 mm outer diameter; World Precision Instruments, Sarasota, FL) and filled with a normal internal solution (in mM): 128 potassium gluconate, 10 NaCl, 1 MgCl_2_, 11 EGTA, 10 HEPES, 3 ATP, and 0.25 GTP (pH was adjusted to 7.3–7.4 with 1N KOH, 290–300 mOsm). Pipette resistances ranged from 3–5 MΩ. In whole cell configuration, access resistance was less than 20 MΩ; access resistance was 80% compensated. For some experiments measuring the ramp current–voltage (I–V) relationship, K^+^-gluconate in the normal internal solution was replaced with Cs^+^-gluconate (pH 7.35 with CsOH), and the extracellular solution contained Na^+^, K^+^, I_h_ (HCN), Ca^2+^, and GABA_A_ channel blockers (in mm: NaCl, 126; 4-aminopyridine, 5; KCl, 2.5; MgCl_2_, 1.2; CsCl, 2; CaCl_2_, 1.4; CoCl_2_, 1; nifedipine, 0.01; HEPES, 20; NaOH, 8; glucose, 10; tetrodotoxin, 0.001; picrotoxin, 0.1). For optogenetic stimulation, a light-induced response was evoked using a light-emitting diode (LED) 470 nm blue light source controlled by a variable 2A driver (ThorLabs, Newton, NJ) with the light path delivered directly through an Olympus 40 water-immersion lens. High fidelity response to light (470 nm) stimulation of Kiss1^ARH^ ::ChR2-mCherry expressing neurons was observed, and both evoked inward currents (in voltage clamp, V_hold_ = −60 mV) or depolarization (in current clamp) were measured. Electrophysiological signals were amplified with an Axopatch 200A and digitized with Digidata 1322A (Molecular Devices, Foster City, CA), and the data were analyzed using p-Clamp software (RRID:SCR_011323, version 9.2, Molecular Devices). The amplitude of the slow EPSP was measured after low pass filtering in order to eliminate the barrage of action potentials riding on the depolarization. The liquid junction potential was corrected for all data analysis.

### Calcium imaging

For calcium imaging, brain slices were placed in a RC-22C slide recording chamber (Harvard/Warner Instruments) and imaged on an inverted Nikon TiE microscope equipped with a Yokogawa CSU-W1 spinning disk confocal head, integrated under NIS Elements v4.20 (Nikon). The preparation, kept at 32°C via a cage incubator (Okolab), was continuously perfused with oxygenated aCSF at a flow rate of 1.25 ml/min. Images were acquired on a Zyla v5.5 sCMOS camera (Andor) at 0.5 Hz. frame-rate, through an 10 x (NA 0.45) or 20 x (NA 0.75) objective, combining 488 nm laser excitation with 500–550 nm emission collection. A single focal plane (z-axis) was maintained using the Nikon Perfect Focus System. Minor tissue drift in the x-y axis was corrected using NIS Elements. Imaging displaying major drift were excluded from final analysis. Changes in Kiss1^ARH^ neuron Ca^2+^ levels were measured in regions of interest (ROIs) comprising the GCaMP6s-positive cell bodies. In all recordings, background fluorescence measured in an ROI drawn on nearby tissue was subtracted from every ROI. [Ca^2+^]_i_ variations after drug applications were assessed as changes in fluorescence signals over baseline (ΔF/F_0_). To normalize the fluorescence value of each cell, we first separated experimental trials into two parts: a baseline period (2 min) corresponding to all the frames recorded before addition of drugs, and a stimulus period, after the onset of the drug (such as bath-applied senktide) application and lasting several minutes. Next, for each ROI we calculated ΔF/F_0_ for each frame (t), where ΔF/F_0_ equals (F_(t)_ – F_0_)/F_0_, and F_0_ was the mean fluorescence value for that ROI for all frames in the baseline period for that trial. The area under the curve (AUC) was calculated over the time periods of 2 min before and 18 min after drug application. Maximal peak reached after drug application was also measured and used in quantitative analysis. Data were averaged across all Kiss1^ARH^ neurons in a slice (two slices per animal), which were used as the statistical unit over a minimum of 3 animals per condition.

### Single cell RT-PCR (scRT-PCR)

Coronal brain sections from the ARH of three female *Stim1*^*kko*^ and three *Kiss1*^*Cre:GFP*^*::Ai32* mice were prepared for electrophysiology and scRT-PCR. The 3-4 slices obtained were divided between electrophysiological recording experiments and single cell harvesting. Single cell dispersion and harvesting was performed as described previously with some modifcations as described below (Bosch et al., 2013; Zhang et al., 2013b). Briefly, the ARH was dissected and digested in papain (7mg/ml in aCSF, Sigma-Aldrich). Gentle trituration using varying sizes of flame polished Pasteur pipets were used to disperse the cells and then they were plated onto a glass bottom dish. A constant flow of oxygenated aCSF (NaCl, 125 mM; KCl, 5 mM; NaH_2_PO_4,_ 1.44 mM; Hepes, 5 mM; D-glucose, 10 mM; NaHCO3, 26 mM; MgSO_4·_7H_2_O, 2 mM; CaCl_2_, 2 mM) was applied to the dish to keep the cells healthy and to clear debris. Fluorescent neurons were visualized under an inverted microscope. The Xenoworks Microinjection system (Sutter Instruments) was used to manipulate a 10 µm tip size glass capillary tube to approach single neurons, apply gently suction and harvest single cells or pools of 10 cells into a siliconized tube containing a solution of 1X Invitrogen Superscript III Buffer (LifeTech), 15U of RNasin (Promega), 10 mM of dithiothreitol (DTT) and diethylpyrocarbonate (DEPC)-treated water in a total of 5 µl for single cells or 8 µl for pools of 10 cells. Corresponding controls were collected at the same time including single neurons (processed without reverse transcriptase) and aCSF from the surrounding area. Hypothalamic tissue RNA was also processed with and without reverse transcriptase. First strand cDNA synthesis was performed on single cells, pools of cells and controls in a 20 µl (single cells) or 25 µl (10 cell pools) volume containing a final concentration of 1X Invitrogen Superscript III Buffer, 30 U of RNasin, 15 mM DTT, 10 mM dNTP, 100 ng Random Primers (Promega), 400 ng Anchored Oligo (dT)_20_ Primer (Invitrogen), 100 U Superscript III Reverse Transcriptase (Life Tech) and DEPC-treated water according to manufactures protocol and stored at −20°C. Clone Manager software (Sci Ed Software) was used to design primers that cross at least one intron-exon boundary. In order to confirm that STIM1 was knocked out, STIM1 primers were designed to include part of exon 2 (see Table 1). Single cell PCR conditions were optimized for primer concentration, magnesium concentration and annealing temperature. Standard curves were generated using hypothalamic cDNA with dilutions from 1:50 to 1:12,800 for primers used for qPCR to determine the efficiency (*E* = 10^(−1/*m*)^-1; table 1). Primer pairs with efficiencies of 90-100% permit the use of the comparative ΔΔCT method for analysis (Livak and Schmittgen, 2001; Pfaffl, 2001).

**Table 1.** Primer Table.

| Gene Name | Accession | Primer | Product | Annealing | <u>Efficiency</u> |  |  |
| --- | --- | --- | --- | --- | --- | --- | --- |
| (encodes for) | Number | Location (nt) | Length (bp) | Temp (°C) | Slope | r<sup>2</sup> | % |
| *Kiss1* (Kiss1)<sup>a,b</sup> | NM_178260 | 64-80 (exon 1) | 120 | 57<sup>a</sup>, 60<sup>b</sup> | -3.410 | 0.989 | 97 |
|  |  | 167-183 (exon 2) |  |  |  |  |  |
| *Stim1* (STIM1)<sup>a</sup> | NM_009287 | 797-816 (exon 2) | 204 | 59 |  |  |  |
|  |  | 981-1000 (exon 3) |  |  |  |  |  |
| *Stim1* (STIM1)<sup>b</sup> | NM_009287 | 821-839 (exon 2) | 135 | 60 | -3.311 | 0.977 | 100 |
|  |  | 937-955 (exon 3) |  |  |  |  |  |
| *Stim2* (STIM2)<sup>a</sup> | NM_001363348 | 620-638 (exon 2) | 172 | 59 |  |  |  |
|  |  | 773-791 (exon 4) |  |  |  |  |  |
| *Stim2* (STIM2)<sup>b</sup> | NM_001363348 | 1784-1803 (exon 11) | 131 | 60 | -3.439 | 0.993 | 95 |
|  |  | 1895-1914 (exon 12) |  |  |  |  |  |
| *Gapdh* (GAPDH)<sup>b</sup> | NM_008084 | 689-706 (exon 4) | 93 | 60 | -3.352 | 0.998 | 99 |
|  |  | 764-781 (exon 5) |  |  |  |  |  |
| *Trpc4* (TRPC4)<sup>a,b</sup> | NM_016984 | 1888-1907 (exon 6) | 116 | 60 | -3.318 | 0.940 | 100 |
|  |  | 1984-2003 (exon 7) |  |  |  |  |  |
| *Trpc5* (TRPC5)<sup>a</sup> | NM_009428 | 2206-2227 (exon 6) | 131 | 63 |  |  |  |
|  |  | 2315-2336 (exon 7) |  |  |  |  |  |
| *Trpc5* (TRPC5)<sup>b</sup> | NM_009428 | 734-753 (exon 2) | 118 | 60 | -3.161 | 0.953 | 100 |
|  |  | 832-851 (exon 3) |  |  |  |  |  |

<sup>a</sup>primers for scRT-PCR.
<sup>b</sup>primers for qPCR.

PCR for *Kiss1, Stim1, Trpc4* and *Trpc5* mRNAs was performed on 3 µl of cDNA from single cells in a 30 µl reaction volume containing 1X GoTaq Flexi buffer (Promega), 2 mM MgCl_2_, 10 mM dNTP, 0.33 µM forward and reverse primers, 2 U GoTaq Flexi Polymerase (Promega) and 0.22 µg TaqStart Antibody (Clontech). 45-50 cycles of amplification were performed on a Bio-Rad C1000 thermocycler and the resulting product visualized with ethidium bromide on a 2% agarose gel.

Quantitative PCR (qPCR) was performed on 3-4 µl of cDNA from pools of 5-10 cells (3-4 pools/animal) in duplicate for the target genes (*Stim1, Stim2, Trpc4 and Trpc5*) and 2 µl in duplicate for the reference gene (*Gapdh*) in a 20 µl reaction volume containing 1X Power SYBR Green PCR Master Mix (Applied Biosystems) and 0.5 µM forward and reverse primers. Forty cycles of amplification were run on a Quant Studio 7 Flex Real-Time PCR System (Applied Biosystems) and the resulting data was analyzed using the comparative ΔΔCT method (Livak and Schmittgen, 2001; Pfaffl, 2001). The relative linear quantity was determined with the 2^-ΔΔCT^ equation (Bosch et al., 2013). The mean of all of the ΔCT values (ΔCT = CT of the target gene – CT of the reference gene) from the controls was used as the calibrator and the data is expressed as fold change in gene expression.

### Drugs

A standard artificial cerebrospinal fluid was used (Qiu et al., 2011). All drugs were purchased from Tocris Bioscience (Minneapolis, MN) unless otherwise specified. Tetrodotoxin (TTX) was purchased from Alomone Labs (Jerusalem, Israel) (1 mM) and dissolved in H_2_O. Thapsigargin (Tg, 2 mM), TacR3 agonist senktide (1 mM) and TRPC4/5 antagonist, HC 070 (from MedChemExpress, 10 mM) were prepared in dimethylsulfoxide (DMSO). Aliquots of the stock solutions were stored as appropriate until needed.

### Data analysis

For qPCR four Kiss1 neuronal pools (10 cells/pool) from each animal were run in duplicate for the mRNAs that encode for STIM1, STIM2 and GAPDH and the mean value of each gene from each animal (n = 3 animals) was used for statistical analysis. Data are expressed as mean ± SEM and were analyzed using an unpaired student’s t-test. In addition, Kiss1 neuronal pools (5-10 cells/pool) were used to determine the expression of *Trpc4* and *Trpc5* in these neurons. For scRT-PCR the number of Kiss1-positive cells harvested from Kiss1^Cre:GFP^ females injected with Cre-dependent ChR2-mCherry or from Kiss1^Cre:GFP^::Ai32 females were used to qualitatively assess the number of Kiss1 neurons with *Stim1, Stim2, Trpc4, Trpc5* and percent expression.

Comparisons between different treatments were performed using a repeated measures, two-way or one-way ANOVA analysis with the *post hoc* Bonferroni’s test. Differences were considered statistically significant if p < 0.05. All data are expressed as mean ± SEM.

## Results

### Validation of conditional deletion of Stim1 in Kiss1 neurons

STIM1 is involved in the regulation of neuronal firing in cerebellar Purkinje neurons (Hartmann et al., 2014; Ryu et al., 2017), dopaminergic neurons (Sun et al., 2017) and hypothalamic arcuate POMC neurons (Qiu et al., 2018b). Initially to see if STIM1 regulates Kiss1^ARH^ neuronal excitability, we measured the mRNA expression of *Stim1* and its close homolog *Stim2* in manually harvested Kiss1 ^ARH^ neurons by quantitative real-time PCR (***Figure 1A***). Based on the qPCR, mRNA levels of *Stim1* were greater than those of *Stim2* in Kiss1 ^ARH^ neurons (***Figure 1A1***). Likewise, in cerebellar Purkinje neurons, *Stim1* is also much more abundant than *Stim2* (Hartmann et al., 2014), while in hippocampal (Berna-Erro et al., 2009) and cortical neurons (Gruszczynska-Biegala et al., 2011) *Stim2* expression levels exceed those of *Stim1*. A qualitative, unbiased sampling of Kiss1^ARH^ neurons (n=60) from ovariectomized *Kiss1*^*Cre*^ females (n =3) revealed that *Stim1* mRNA was expressed in 81.7 ± 7.6 percent and *Stim2* mRNA was detected in 81.2 ± 2.7 percent of Kiss1^ARH^ neurons with 70 percent of neurons expressing both *Stim1* and *Stim2*.

**Figure 1.**
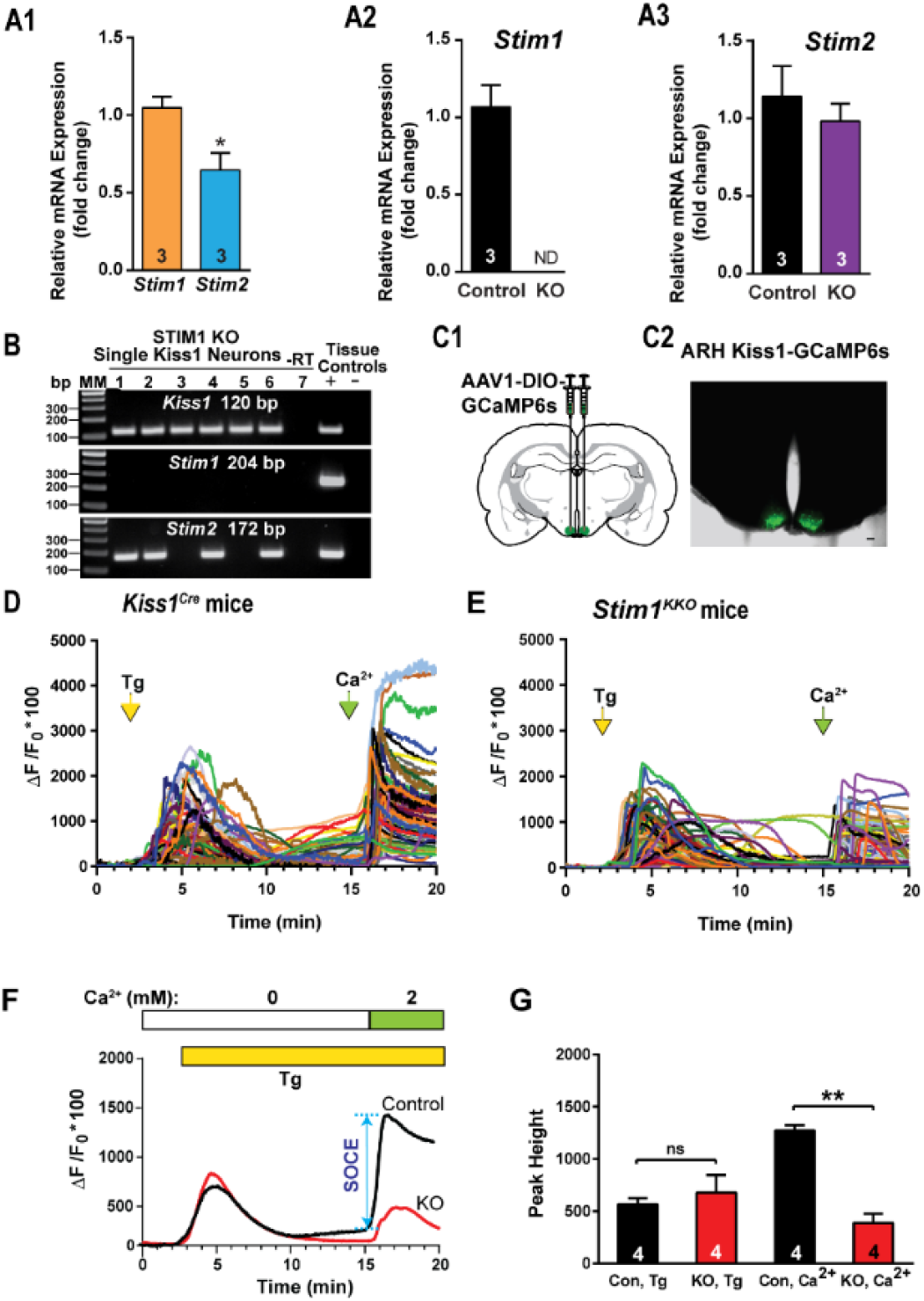
Expression Patterns of *Stim1 and 2* in the arcuate Kiss1 neurons. **A1-A3**, quantitative PCR assay measuring *Stim1* and *Stim2* in Kiss1^ARH^ neuronal pools (n = 3 animals, 10 cells in each pool, 4 pools/animal) from *Kiss1*^*Cre*^ control and *Stim1*^*kko*^ female mice (n=3 animals per group). **A1**, comparison between *Stim1* and *Stim2* in controls only. Bar graphs represent mean ± SEM (Unpaired *t*-test, *t*_(4)_ = 3.079, * *p* = 0.0370); **A2**, *Stim1* was non-detectable (ND) in the STIM1^KKO^ neuronal pools (Unpaired *t*-test, *t*_(4)_ = 7.559, ** *p* = 0.0016); **A3**, the *Stim2* expression level of Kiss1^ARH^ neurons was not different between *Kiss1*^*Cre*^ control and *Stim1*^*kko*^ female mice (Unpaired *t*-test, *t*_(4)_ = 0.7143, *p* = 0.5145). **B**, representative gels illustrating mRNA expression of *Stim1* and *Stim2* in single Kiss1^ARH^ neurons from *Stim1*^kko^ mice. The expected base pair (bp) sizes are *Kiss1*, 120 bp; Stim1, 204 bp; Stim2, 172 bp. A single neuron was processed without reverse transcriptase (-RT) and RNA extracted from hypothalamic tissue was used as positive (+, with RT) and negative (-, without RT) tissue controls. MM, molecular marker. **C**, left, schematic of a coronal section showing the bilateral viral injections in the ARH with AAV-DIO-GCaMP6s. Right, photomicrograph showing a coronal section confirming targeted bilateral injections of DIO-GCaMP6s into the ARH. **D** and **E**, representative traces of GCaMP6s activity based on cytosolic Ca^2+^ measurements in Kiss1^ARH^ neurons from *Kiss1*^*Cre*^*:GCaMP6s* mice (D) and *Stim1*^*kko*^*:GCaMP6s* mice (E). ER Ca^2+^ stores were depleted with 2 µM thapsgargin, a SERCA inhibitor, after 20 min of perfusion with aCSF containing 0 mM Ca^2+^. SOCE was evaluated by substituting the extracellular aCSF containing 0 mM Ca^2+^ with aCSF containing 2 mM Ca^2+^. **F**, averaged traces from D and E revealed that deletion of *Stim1* in Kiss1^ARH^ neurons attenuated the store-operated Ca^2+^ entry (SOCE). **G**, bar graphs summarizing the effects of depletion of Ca^2+^ store by Tg and Ca^2+^ influx (SOCE) in Kiss1^ARH^ neurons from Kiss1Cre:GCaMP6s and Stim1^kko^:GCaMP6s mice (two-way ANOVA: main effect of treatment (F_(1,3)_ = 13.84, *p* = 0.0338), main effect of time (F_(1,3)_ = 5.199, p = 0.1069) and interaction (F_(1,3)_ = 52.14, *p* = 0.0055); n = number of slices; *post hoc* Bonferroni test, \*\**p* < 0.01, for SOCE; ns = no significant, for depletion of Ca^2+^ store.

To elucidate the functional role of STIM1 in Kiss1 neurons, we generated mice that lack STIM1 selectively in Kiss1 neurons (*Stim1*^*kko*^, detailed in Methods). We confirmed the *Stim1* deletion in *Stim1*^*kko*^ mice using single cell quantitative PCR of pools of harvested Kiss1^ARH^ neurons (n= 3 animals) (***Figure 1A2***). Consistent with the scRT-PCR results (***Figure 1B***), *Stim1* mRNA was undetectable in *Stim1*^*kko*^ neurons (***Figure 1A2***), whereas there was no reduction in *Stim2* mRNA expression (***Figure 1A3***). In contrast, *Stim1* mRNA was still expressed in the majority of adjacent nonfluorescent neurons obtained from both *Stim1*^*kko*^ and *Kiss1*^*Cre*^ mice.

### Stim1 deletion reduces Store Operated Calcium Entry (SOCE)

SOCE constitutes an important source of calcium entry and signaling in neurons. Depletion of ER Ca^2+^ stores causes the ER Ca^2+^ sensor STIM proteins (STIM1 and STIM2) to interact with and activate cell surface Ca^2+^ release-activated Ca^2+^ (CRAC) channels, thereby resulting in a second wave of cytoplasmic Ca^2+^ rise (Moccia et al., 2015). Genetic suppression of *Stim1* in neural progenitor cells results in abrogation of this second wave of calcium rise that constitutes SOCE (Somasundaram et al., 2014). We asked whether deletion of *Stim1* in Kiss1^ARH^ neurons (*Stim1*^*kko*^) attenuates neuronal SOCE. Kiss1^Cre^ and Stim1^kko^ mice received bilateral ARH injections of GCaMP6 viral vector (***Figure 1C1, C2***), and the Kiss1^ARH^ or *Stim1*^*kko*^ neurons with GCaMP6s in slices were imaged using spinning disk confocal microscopy (***Figure 2-video supplement 1***). ER Ca^2+^ stores were released by treatment with 2 μM thapsigargin (Tg), a blocker of the SERCA (sarcoplasmic/endoplasmic reticulum Ca^2+^ ATPase) pump. As expected, Tg treatment of neurons bathed in Ca^2+^-free aCSF generated an initial wave of cytoplasmic Ca^2+^ release ([Ca^2+^]_i_) as measured by an increase in GCaMP6s activity both in control and *Stim1*-deleted neurons (***Figure 1D, E and F***). As long as neurons were kept in Ca^2+^-free aCSF, the ER stores remained empty, a situation that was presumably sensed by the Ca^2+^ sensor STIMs. Upon switching to a normal aCSF containing 2 mM Ca^2+^, an immediate SOCE response was observed as a second wave of cytoplasmic Ca^2+^ rise. Consistent with a role for STIM1 regulation, we observed an attenuation of SOCE in *Stim1*^*kko*^ neurons (***Figure 1D, E, F and G***: ΔF/F_0_*100 =1274.5 ± 49.4, n = 4, Kiss1^ARH^ group versus 389.0 ± 86.1, n = 4, Stim1^kko^ group, which was measured from the 15 minute time point to the peak, unpaired t-test, *t*_(6)_ = 8.921, *p* = 0.0001, \*\*\**p* < 0.005), indicating that STIM1 plays a major role in SOCE after Tg-induced ER Ca^2+^ depletion in Kiss1^ARH^ neurons as has been shown in other CNS neurons (Guner et al., 2017; Pavez et al., 2019).

**Figure 2.**
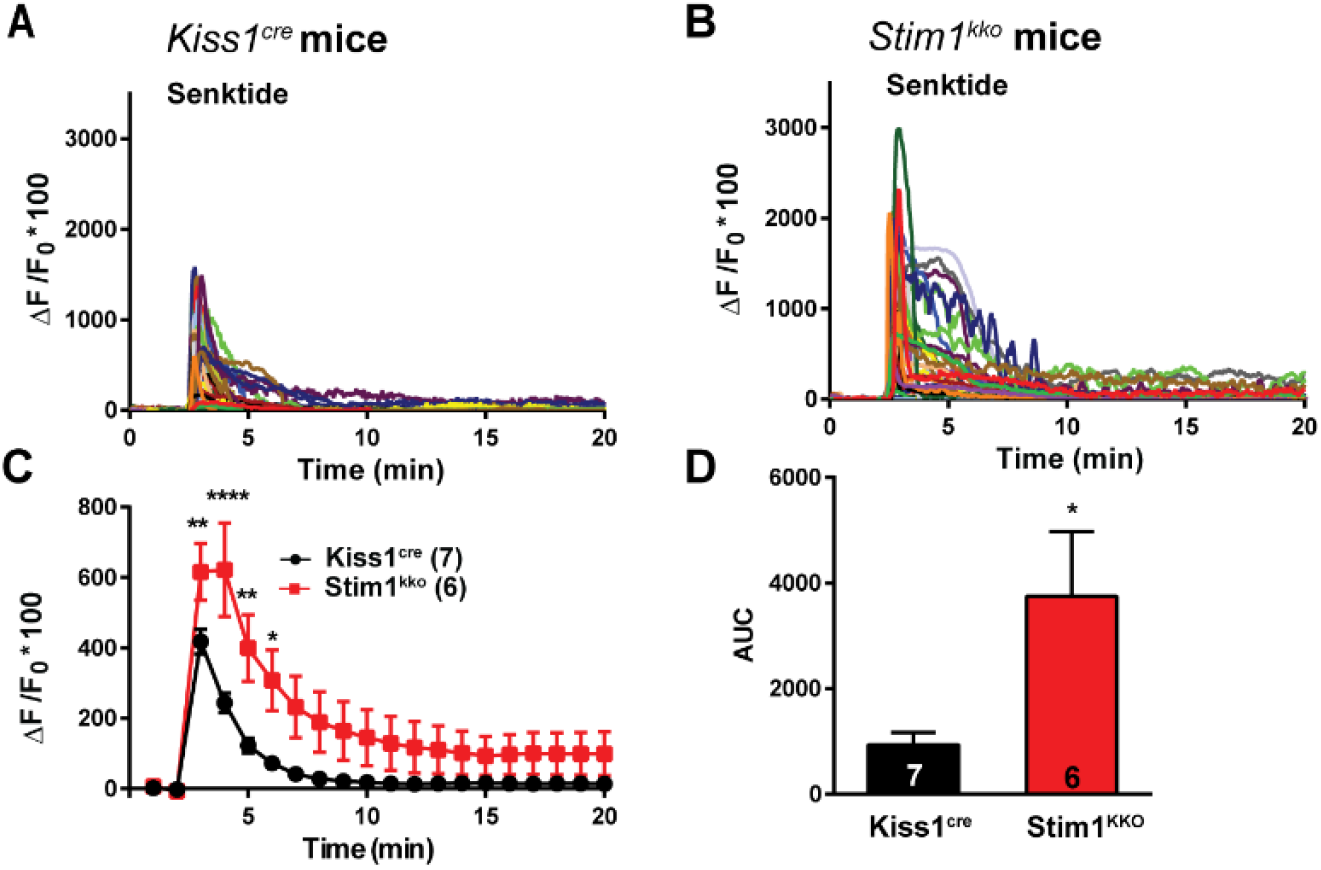
Senktide-induced increase in [Ca^2+^]_i_ is augmented by deletion of *Stim1* in GCaMP6s-expressing Kiss1^ARH^ neurons from *Kiss1*^*Cre*^ and *Stim1*^*kko*^ mice. **A** and **B**, representative traces of senktide-induced [Ca^2+^]_i_in Kiss1^ARH^ neurons from *Kiss1*^*Cre*^ (A) and *Stim1*^*kko*^ (B) mice. Traces represent individual cells within a single slice. **C**, summary of the potentiation of senktide-induced [Ca^2+^]_i_ by deletion of *Stim1*. Two-way ANOVA: main effect of treatment (F_(1,11)_ = 5.265, *p* = 0.0424), main effect of time (F_(19,209)_ = 42.69, p < 0.0001) and interaction (F_(19,209)_ = 6.486, *p* < 0.0001); n = number of slices; *post hoc* Bonferroni test, \*\*\*\**p* < 0.001; \*\**p* < 0.01; \**p* < 0.05. **D**, AUC of Kiss1^ARH^ neurons from *Kiss1*^*Cre*^ and *Stim1*^*kko*^ mice from C. There was a significant difference (Unpaired *t*-test, *t*_(11)_ = 2.430, \**p* = 0.0334) between the two groups.

### TacR3 -induced increase in [Ca^2+^]_i_ is augmented by deletion of Stim1

TacR3 classically couples to a Gαq protein-calcium signaling and excites Kiss1^ARH^ neurons (de Croft et al., 2013; Ruka et al., 2013; Qiu et al., 2016). Calcium is of critical importance to neurons as it participates in the transmission of depolarizing signals and contributes to synaptic activity (Brini et al., 2014). Therefore, we tested whether STIM1 can modulate TacR3-mediated calcium responses. We first measured the effects of the TacR3, which is Gq-coupled, agonist senktide on GCaMP6s-expressing Kiss1^ARH^ neurons in arcuate slices from *Kiss1*^*Cre*^ mice; senktide (1 μM) rapidly induced an increase in [Ca^2+^]_i_ (***Figures 2A,C***). Next, we investigated if STIM1 contributes to intracellular rise in [Ca^2+^]_i_ after senktide activation. Indeed, deletion of *Stim1* significantly augmented the peak TacR3-mediated response by ∼three-fold (ΔF/F_0_*100 = 244.0 ± 27.7, n = 7 slices, Kiss1^ARH^ group versus 622.1 ± 133.2, n = 6 slices), *Stim1*^*kko*^ group; two-way ANOVA: main effect of treatment (F_(1,11)_ = 5.265, *p* = 0.0424), main effect of time (F_(19,209)_ = 42.69, p < 0.0001) and interaction (F_(19,209)_ = 6.486, *p* < 0.0001); *post hoc* Bonferroni test, \*\*\*\**p* < 0.001; \*\**p* < 0.01; \**p* < 0.05). (***Figures 2B, C***). Likewise, the area under the curve was significantly increased in the *Stim1*^*kko*^ group by four-fold (Kiss1^cre^: 954.8 ± 200.4, n = 7 versus Kiss1^kko^: 3746.0 ± 1227.0, n = 6) (***Figure 2D***).

### Deletion of STIM1 enhances slow EPSP in Kiss1^ARH^ neurons

Kiss1^ARH^ neurons are the chief component of the GnRH pulse generator circuit (Navarro et al., 2009; Lehman et al., 2010; Navarro et al., 2011; Okamura et al., 2013), such that they synchronize their activity to trigger the release of peptides to drive pulsatile release of GnRH (Qiu et al., 2016; Clarkson et al., 2017). To investigate if STIM1 modulates the activity of Kiss1^ARH^ neurons, we bilaterally injected AAV1-Ef1a-DIO-ChR2:mCherry into the arcuate nucleus of *Kiss1*^*Cre*^ and *Stim1*^*kko*^ mice. To verify that *Trpc5* mRNAs is co-localized in these Kiss1^ARH^ neurons, we harvested 50 Kiss1^ARH^ neurons from 2 females and did scRT-PCR for *Trpc5* and *Trpc4* expression. The single-cell analysis revealed that *Trpc5* transcript was detectable in 82% of Kiss1^ARH^ neurons, but *Trpc4* mRNA was not detected in Kiss1^ARH^ neurons (***Figure 3A***). Moreover, quantitative single cell PCR documented that *Trpc5* but not *Trpc4* mRNA was expressed in Kiss1^ARH^ neurons (***Figure 3B***). With whole-cell recording we verified the expression of TRPC5 channels by documenting the senktide - induced typical double-rectifying I/V plot characteristic of the activation of TRPC5 channels (***Figure 3C***) as we previously reported (Kelly et al., 2018). Initially, whole-cell patch recording in *Kiss1*^ARH^ neurons from ovariectomized *Kiss1* or *Stim1*^*kko*^ female mice revealed that there was no difference in the resting membrane potential (RMP: *Kiss1*: −66.0 ± 1.7 mV, n = 38 versus *Stim1*^kko^: −68.2 ± 0.9 mV, n = 58) or membrane capacitance (C_m_: *Kiss1*: 25.0 ± 1.0 pF, n = 38 versus *Stim1*^kko^: 27.3 ± 0.9 pF, n = 58). However, there was a significant difference in the membrane input resistance (R_in_: *Kiss1*: 524.2 ± 42.4 Ω, n = 38, versus *Stim1*^kko^: 417.3 ± 26.9 Ω, n = 58, unpaired two-tailed *t* test, t_(94)_ = 2.242, p = 0.0273), which has also been reported with *Stim1* knockout in cerebellar Purkinje neurons (Ryu et al., 2017). Kiss1^ARH^ neurons expressing ChR2-mCherry in slices were photostimulated at 20 Hz for 10 s (***Figure 3-video supplement 1***) to generate slow EPSPs as previously described (Qiu et al., 2016). As we had hypothesized, deletion of *Stim1* augmented the slow EPSP induced by high-frequency optogenetic stimulation (***Figure 3D-F***). Also in the presence of TTX to block voltage-gated Na^+^ channels, we observed that senktide induced larger inward currents in Kiss1^ARH^ neurons from *Stim1*^*kko*^ mice versus *Kiss1*^*Cre*^ mice (***Figure 4A-C***). Although the senktide-induced cation current was significantly increased by *Stim1* deletion, the I/V plots revealed that the reversal potential for the current was not different between Kiss1^ARH^ neurons recorded from *Stim1*^*kko*^ mice or *Kiss1*^*Cre*^ mice (*Kiss1*: −10.5 ± 2.1 mV, n = 4, vs *Stim1*^kko^: −9.8 ± 2.2 mV, n = 4, unpaired two-tailed *t* test, t_(6)_ = 0.2503, p = 0.8107) (***Figure 4D-F***). These results indicate that STIM1 expression governs the activity of TRPC5 channels, which contribute to the synchronous activity of Kiss1^ARH^ neurons.

**Figure 3.**
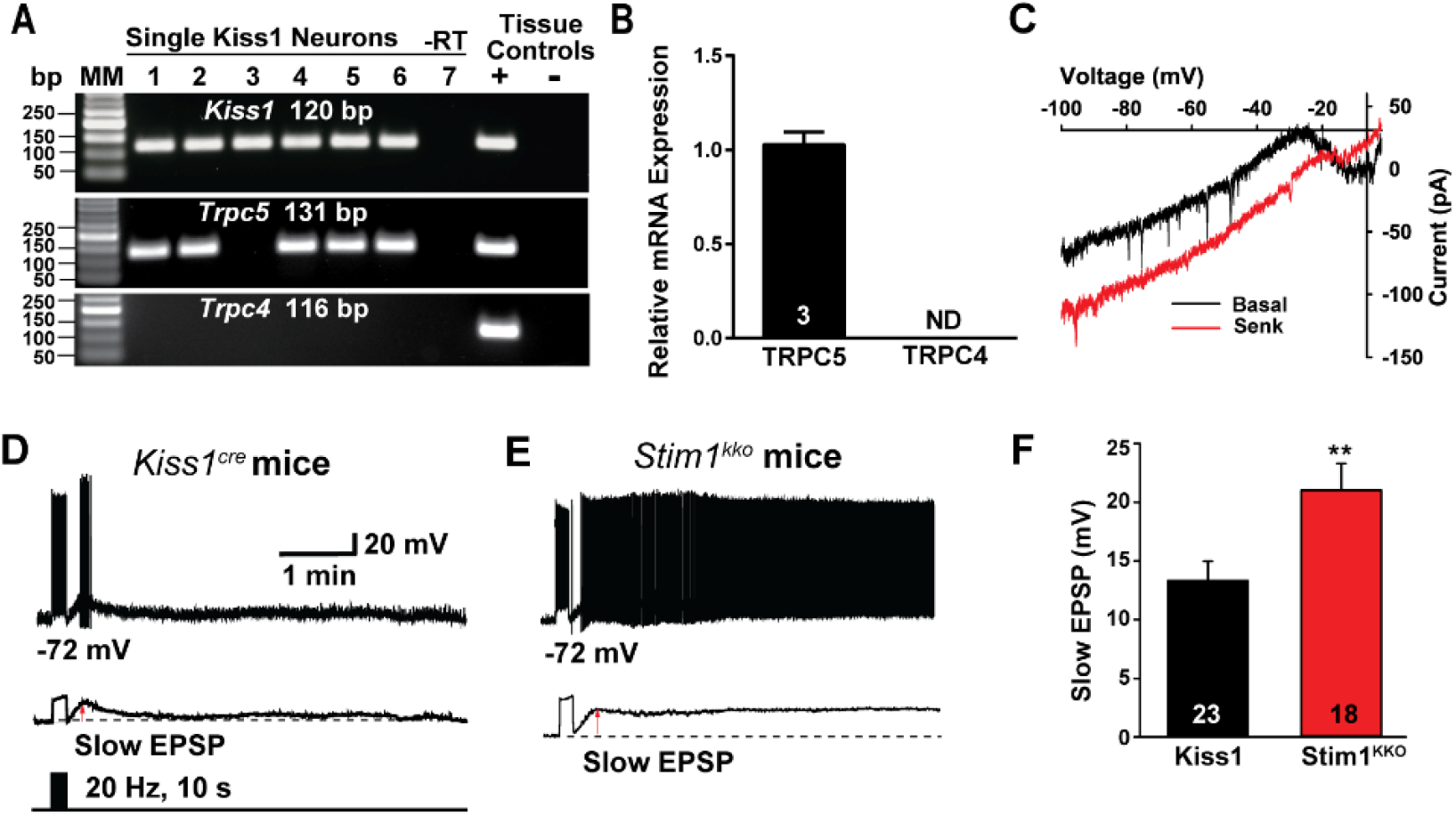
Deletion of *Stim1* augments high-frequency optogenetic stimulation-induced slow EPSP depolarization in Kiss1^ARH^ neurons. **A**, representative gel image illustrating the mRNA expression of *Trpc5* channel subunit in Kiss1^ARH^ neurons harvested from female mice. The expected size of PCR products for *Kiss1* and *Trpc5* are indicated. *Trpc4* mRNA was not detected in Kiss1^ARH^ neurons. MM is the molecular marker; –RT indicates a harvested Kiss1 neuron reacted without RT; + indicates positive tissue control (with RT); – indicates negative tissue control (without RT) using cDNA from mouse medial basal hypothalamic tissue; RT, reverse transcriptase. **B**, quantitative single cell PCR (3 x 10 cell pools per animal, n = 3 animals) verified that *Trpc5* was expressed in Kiss1^ARH^ neurons, whereas *Trpc4* mRNA was not detected (Unpaired *t*-test for the left, *t*_(4)_ = 15.67, **** p < 0.0001). **C**, the I-V relationship for the Senk-induced inward current recorded in Kiss1^ARH^ neurons from *Kiss1*^*Cre*^ mice using a Cs^+^ internal solution (n= 4) revealed a reversal of −10 mV and rectification. **D, E**, high-frequency optogenetic stimulation (20 Hz, 10 s) generated slow EPSPs in a ChR2-expressing Kiss1^ARH^ neuron from control *Kiss1* mice (**D**) and in a ChR2-expressing Kiss1^ARH^ neuron from *Stim1*^*kko*^ mice (**E**). The lower trace shows the slow EPSP after low-pass filtering from D and E (arrow), respectively. **F**, summary of the effects of *Stim1* deletion on the slow EPSP amplitude. Bar graphs represent the mean ± SEM (Unpaired *t*-test, *t*_(39)_ = 2.802, \*\**p* = 0.0079).

**Figure 4.**
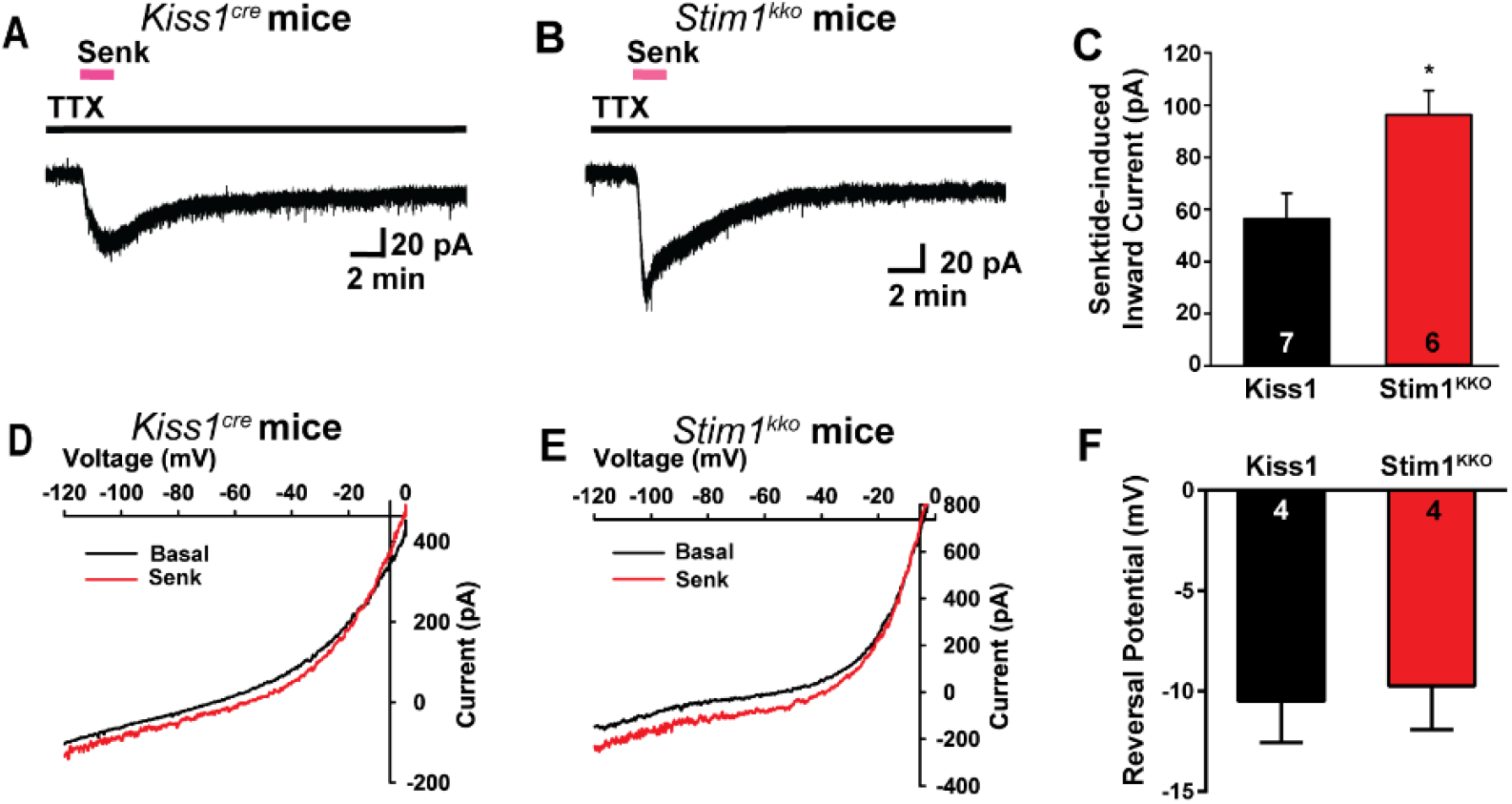
Deletion of *Stim1* augments senktide-induced depolarization in Kiss1^ARH^ neurons. **A** and **B**, rapid bath application of senktide (1 µM) induced an inward current in the presence of fast sodium channel blockade (TTX, 1µM) in Kiss1^ARH^ neurons from *Kiss1*^*Cre*^ and *Stim1*^*kko*^ mice. V_hold_ = - 60 mV. **C**, summary of the effects of senktide in Kiss1^ARH^ neurons from *Kiss1*^*Cre*^ and *Stim1*^*kko*^ mice (Unpaired *t*-test, *t*_(11)_ = 2.929, \**p* = 0.0137). Data points represent the mean ± SEM. Cell numbers are indicated. **D** and **E**, the I-V relationship before and during the peak response of senktide (Senk) in Kiss1^ARH^ neurons from *Kiss1*^*Cre*^ (D) and *Stim1*^*kko*^ (E) mice indicated that the reversal potential of the nonselective cation current was ∼ −10 mV. **F**, summary of the reversal potentials of the senktide-induced cation current recorded in Kiss1^ARH^ neurons from *Kiss1*^*Cre*^ and *Stim1*^*kko*^ mice. Bar graphs represent the mean ± SEM (unpaired two-tailed *t* test, t_(6)_ = 0.2503, p = 0.8107).

### NKB agonist activates TRPC5 channels in Kiss1^ARH^ neurons from Kiss1^Cre^ and Stim1^kko^ mice

Based on our previous findings that TPRC5 channel protein is expressed in Kiss1^ARH^ neurons and is activated by the NKB agonist senktide (Qiu et al., 2011; Kelly et al., 2018), we investigated the contribution of TRPC5 channels to generating the slow EPSP. We used a ratio method in which a slow EPSP was generated by optogenetic stimulation (20 Hz, 10 s) of Kiss1^Cre^:ChR2 neurons and then stimulated again 10 min later after drug exposure (Qiu et al., 2016). For the Kiss1^ARH^ neurons from ovariectomized female *Kiss1:Ai32* mice, the RMP, C_m_ and R_in_ were −72.7 ± 0.8 mV, 22.4 ± 0.7 pF and 458.6 ± 23.3 Ω, n = 50, respectively. Using this protocol, we found that the slow EPSP was inhibited by perfusing the TRPC4/5 channel blocker HC 070 (100 nM) (Just et al., 2018) for 5 minutes, and the ratio was significantly decreased from 60 to 30 percent (***Figure 5A-C***). Since *Trpc4* mRNA is not expressed in Kiss1^ARH^ neurons (**Figure 3A, B**), we would argue that TRPC5 channels mediate the slow EPSP in these neurons. To elucidate the TRPC5 channel contribution to the postsynaptic activity of Kiss1^ARH^ neurons from *Stim1*^*kko*^ mice, we perfused TTX to block fast sodium channels and found that HC 070 significantly suppressed the senktide-induced inward current (***Figure 5D, E and F***).

**Figure 5.**
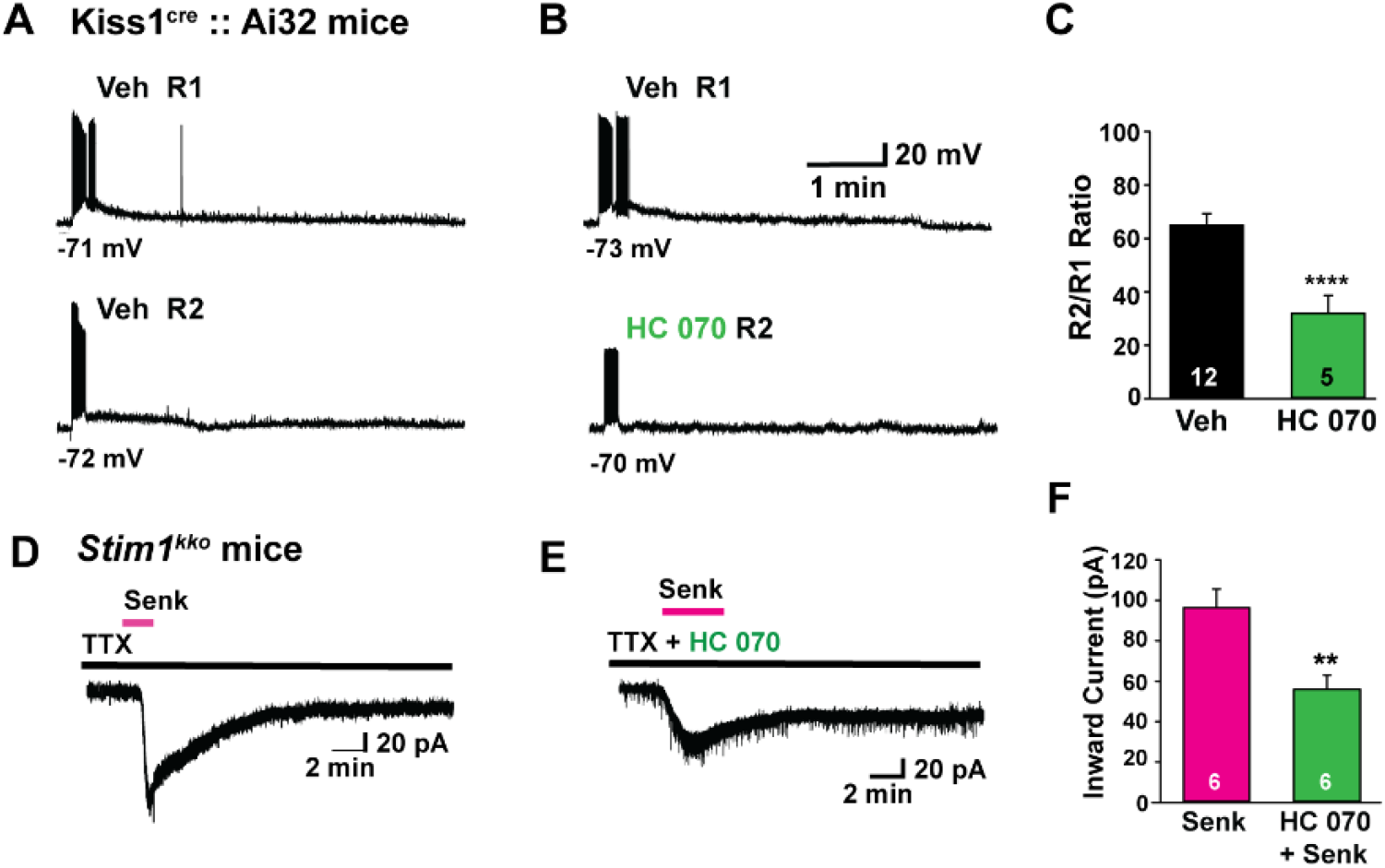
Stim1 deletion augments senktide-induced Kiss1^ARH^ neuronal excitability through TRPC channel activation. **A**-**C**, high frequency photo-stimulation – induced slow EPSP in Kiss1^ARH^ neurons from *Kiss1*^*Cre*^*::Ai32* mice is antagonized by TRPC5 channel blocker. **A**–**B**, representative traces of high-frequency optogenetic stimulation-induced slow EPSPs in the absence (A) or presence (B) of TRPC4/5 channel blocker HC 070 (100 nM). **C**, summary of the effects of HC 070 on the slow EPSP (Un-paired t-test, *t*_(15)_ = 4.122, \*\*\*\**p* = 0.0009). **D-F**, *Stim1* deletion augments senktide-induced inward current, which is antagonized by the TRPC5 channel blocker. **D**–**E**, representative traces of senktide (1 μM)-induced inward current in *Stim1*^*kko*^ neurons perfused with TTX (1 µM) in the absence (D) or presence (E) of TRPC4/5 blocker HC 070 (100 nM). **F**, summary of the effects of HC 070 on the senktide-induced inward current (Un-paired t-test, *t*_(10)_ = 3.457, \*\**p* = 0.0062). Data points represent the mean ± SEM. Cell numbers are indicated.

### Stim1 deletion in Kiss1^ARH^ neurons has minimal effects on estrous cycle

*Stim1*^*kko*^ mice on the C57BL/6 background were viable at the expected Mendelian ratio and did not show any difference in the time to vaginal opening (Stim1^kko^ mice: postnatal day 30.2 ± 0.8, n = 21 versus Kiss1^Cre^mice: postnatal day 29.1 ± 0.8, n = 19, Unpaired *t* test, *t*_(38)_ = 1.003, *p* = 0.3222). However, since kisspeptin neurons are responsible for the maintenance of the reproductive cycle (Seminara et al., 2003; d’Anglemont de Tassigny et al., 2007; Mayer et al., 2010), and *Stim1* deletion facilitated the synchronous firing of Kiss1^ARH^ neurons, we measured the effects of *Stim1* deletion in Kiss1 neurons on the estrous cycle. We monitored the estrous cycle of *Stim1*^*kko*^ and *Kiss1*^*Cre*^ female mice with vaginal lavage for two weeks before ovariectomy for the metabolic studies (see below). *Stim1*^*kko*^ female mice exhibited prolonged estrous cycles versus the *Kiss1*^*Cre*^ females (***Figure 6A,B,C versus 6D,E,F*)** with a slight prolongation of estrous days (***Figure 6H***). Although more in depth analysis is warranted (*i*.*e*., measurement of pulsatile LH), the results were not unexpected since augmented synchronous activity of Kiss1^ARH^ neurons, as we documented at the cellular level, should still drive luteinizing hormone (LH) pulses in these female mice (Qiu et al., 2016; Clarkson et al., 2017).

**Figure 6.**
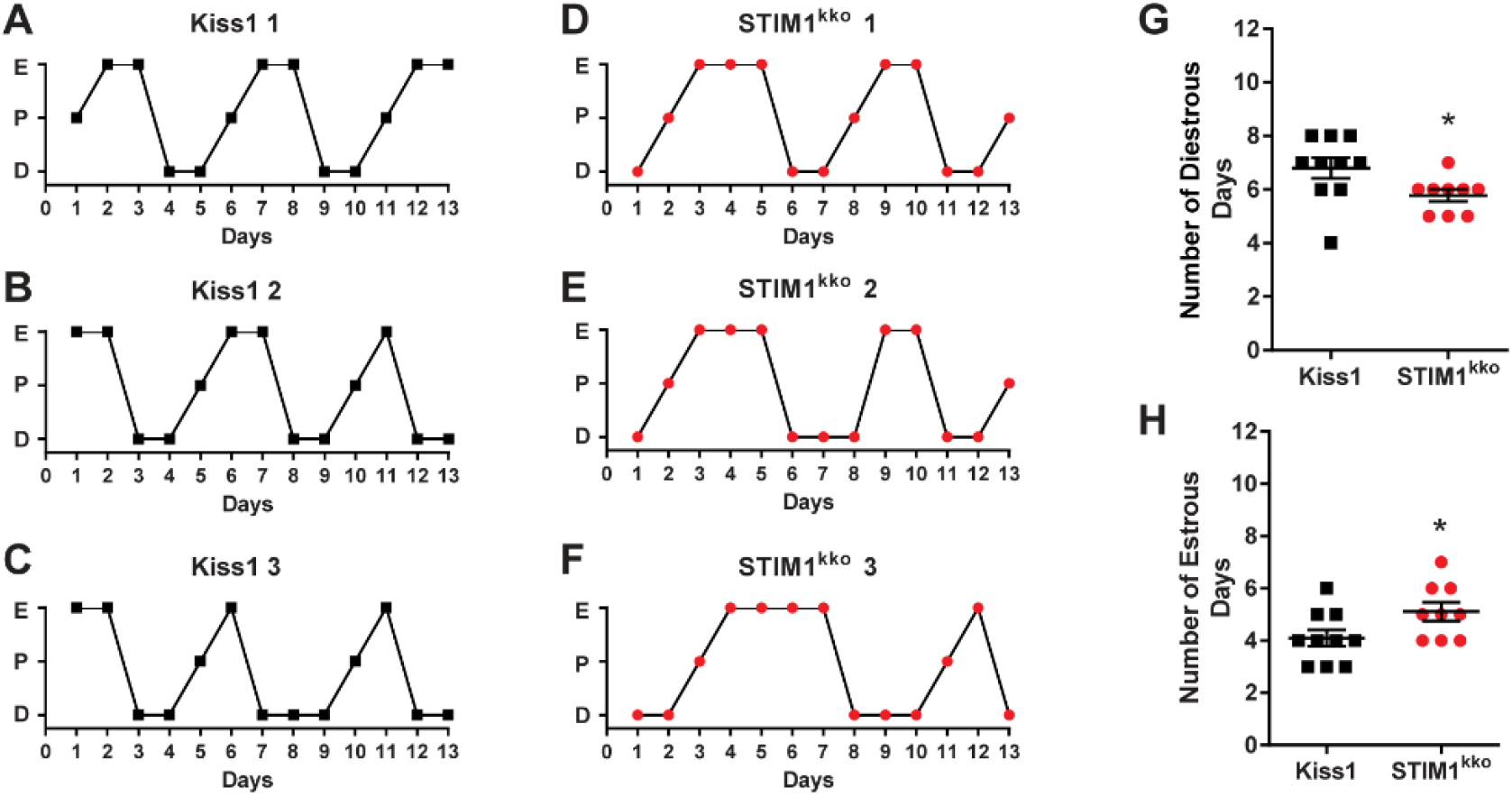
*Stim1*^*kko*^ mice exhibit more estrous days. **A-F**, representative estrous cycle data from three representative control *Kiss1*^*Cre*^ and three *Stim1* ^*kko*^ mice over a thirteen-day period. Vaginal lavage was done daily at 0930 h, and cell cytology was observed and recorded as Diestrus (D), Proestrus (P) or Estrus (E). Summary data for the number of Diestrous days (**G**) and Estrous days (**H**) during the 13 day period was compared between *Kiss1*^*Cre*^ (n = 10) and *Stim1* ^*kko*^ mice (n = 9) (unpaired, two-tailed t test for G, *t*_(17)_ = 2.215, *p = 0.0407; unpaired two-tailed t-test for H, *t*_(17)_ = 2.151, *p = 0.0461).

### Stim1 deletion in Kiss1^ARH^ neurons protects ovariectomized females against diet-induced obesity

Subsequently, the same two cohorts of female mice, *Stim1*^*kko*^ (n=10) and the littermate control *Kiss1*^*Cre*^ (n=10) mice, were ovariectomized at around 3 months of age and put on a high fat diet for eight weeks (see Methods). Over this time period, there was significantly less gain in body weight in the *Stim1*^*kko*^ versus the *Kiss1*^*Cre*^ mice (***Figure 7A, B***). Moreover, the average fat mass of *Stim1*^*kko*^ mice was significantly lighter than that of *Kiss1*^*Cre*^ controls by week 6 (*Stim1*^*kko*^ versus *Kiss1*^*Cre*^ mice fat mass: 7.6 ± 0.9 g, n=10 versus 11.4 ± 1.1 g, n=10) (***Figure 7C***). The lean mass of *Stim1*^*kko*^ mice was also significantly less versus the *Kiss1*^*Cre*^ mice (*Stim1*^*kko*^ versus the *Kiss1*^*Cre*^ mice lean mass: 16.9 ± 0.4 g, n=10 versus 18.9 ± 0.4 g, n=10) (***Figure 7D***). After 6 weeks, both *Stim1*^*kko*^ and *Kiss1*^*Cre*^ controls were assessed for glucose tolerance using an *i*.*p*. glucose tolerance test (see Methods). Both *Stim1*^*kko*^ and *Kiss1*^*Cre*^ females started at relatively the same blood glucose levels after an overnight fast (***Figure 7E***, *time 0*), suggesting similar whole-body homeostatic conditions after fasting. However, *Stim1*^*kko*^ female mice had significantly lower glucose levels after *i*.*p*. glucose compared to *Kiss1*^*Cre*^ females, indicating that *Stim1*^*kko*^ females were more glucose tolerant compared to *Kiss1*^*Cre*^ controls. *Stim1*^*kko*^ females had a significantly higher glucose clearance rate than controls based on the integrated area under the curve (*Stim1*^*kko*^ versus the *Kiss1*^*Cre*^ controls AUC: 20,232 ± 868 mg/dL × min, n = 6 versus 22,622 ± 624 mg/dL × min, n = 6). Finally, when both groups were euthanized after eight weeks on HFD and the tissues harvested, both the intrascapular brown adipose tissue (iBAT) and perigonadal adipose tissue (GAT) were dissected from each mouse and weighed. Both iBAT and GAT masses were significantly lighter in the *Stim1*^*kko*^ versus the *Kiss1*^*Cre*^ females (*Stim1*^*kko*^ versus the *Kiss1*^*Cre*^ iBAT: 73.3 ± 6.0 mg, n=10 versus 97.3 ± 9.6 mg, n=10; *Stim1*^*kko*^ versus the *Kiss1*^*Cre*^ GAT: 1.5 ± 0.2 g, n=10 versus 2.3 ± 0.2 g, n=10) (***Figures 7F, G***). Overall, these results suggest that conditional deletion of *Stim1* in Kiss1^ARH^ neurons affords some protection against diet-induced obesity. However, we cannot overlook the possibility that deletion of *Stim1* in Kiss1-expressing hepatocytes might contribute to this metabolic phenotype (Song et al., 2014).

**Figure 7.**
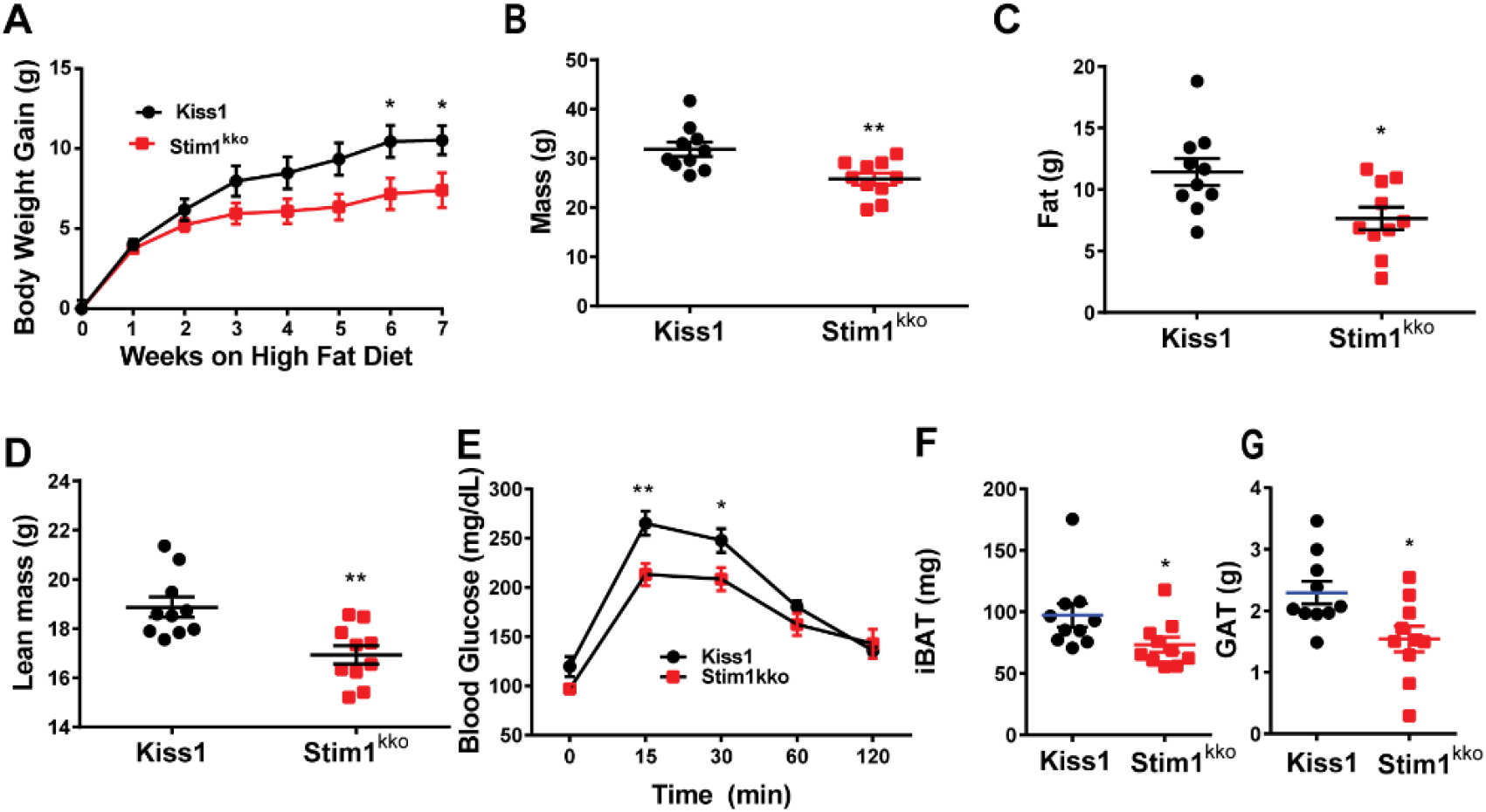
Ablation of Stim1 in Kiss1 neurons attenuates body mass, fat, and lean in mice on a high fat diet. *Stim1*^*kko*^ and *Kiss1*^*Cre*^ littermate control females were ovariectomized and fed a high fat diet (HFD; 45% kcal from fat) for seven weeks. **A**, body-weight gain measured once a week for seven weeks. The high fat diet caused significant weight gain in both groups relative to their baseline with the *Kiss1*^*Cre*^ females gaining significantly more weight by 6 weeks [two-way ANOVA: main effect of treatment (F_(1, 18)_ = 3.839, p = 0.0657), main effect of time (F_(7, 126)_ = 98.07, p < 0.0001) and interaction (F_(7, 126)_ = 4.645, p = 0.0001); Kiss1 control, n = 10, Stim1^kko^, n = 10; *post hoc* Bonferroni test, *p < 0.05]. **B-D**, mass (B), total body fat (C) and lean mass (D) measured by an EchoMRI Whole Body Composition Analyzer. Lean mass did not include bone and fluids within organs. The difference in mass (B), body fat (C) and lean mass (D) between the groups was significantly different by 6 weeks on high fat diet (unpaired, two-tailed t-test for B, *t*_(18)_ = 3.222, **p = 0.0047; unpaired two-tailed t test for C, *t*_(18)_ = 2.662, *p = 0.0159; unpaired, two-tailed t test for D, *t*_(18)_ = 3.489, *p = 0.0026). **E**, six weeks after high fat diet, there was a significant difference in GTTs between the two groups (two-way ANOVA: main effect of treatment (F_(1, 9)_ = 6.282, p = 0.0335), main effect of time (F_(4, 36)_ = 88.01, p < 0.0001) and interaction (F_(4, 36)_ = 3.527, p = 0.0158); *Kiss1*^*Cre*^, n = 6, *Stim1*^*kko*^, n = 5; *post hoc* Bonferroni test, **p < 0.01, *p < 0.05). **F** and **G**, both interscapular brown adipose tissue (iBAT) and perigonadal adipose tissue (GAT) mass of *Stim1*^*kko*^ were lighter than that of *Kiss1*^*Cre*^ mice on a fat diet after eight weeks (unpaired, two-tailed t test for iBAT, *t*_(18)_ = 2.127, *p = 0.0475; unpaired two-tailed t-test for GAT, *t*_(18)_ = 2.711, *p = 0.0143).

## Discussion

For the first time, we show that conditional knockout of *Stim1* significantly reduces store-operated Ca^2+^ entry (SOCE) in Kiss1^ARH^ neurons following thapsigargin-mediated depletion of Ca^2+^ stores. Based on single cell qPCR analysis, *Stim1* mRNA was expressed at approximately two-fold higher levels than *Stim2* in Kiss1^ARH^ neurons, and deletion of *Stim1* did not alter expression of *Stim2* in Kiss1^ARH^ neurons—*i*.*e*., there was no developmental compensation. Selective deletion of *Stim1* in Kiss1^ARH^ neurons augmented the TacR3-mediated increase in [Ca^2+^]_i_ and synchronous activity of Kiss1^ARH^ neurons by almost 4-fold. Whole-cell recording revealed that the slow EPSP induced by high-frequency optogenetic stimulation of Kiss1^ARH^:ChR2 neurons was significantly enhanced by *Stim1* deletion. This augmentation of the slow EPSP was mediated by TacR3 coupling to TRPC 5 channel activation since the senktide-induced inward current was equally enhanced. Moreover, the inward current exhibited the signature double rectifying I/V plot of TRPC 5 channels and was antagonized by the TRPC 4/5 channel blocker HC070. The enhanced TacR3 signaling in *Stim1*^*kko*^ female mice afforded some protection against diet-induced obesity and glucose intolerance.

Mammalian TRPC channels can be activated by G protein-coupled receptors and receptor tyrosine kinases (Clapham, 2003; Ambudkar and Ong, 2007) and are one of the major targets for group I metabotropic glutamate receptor (mGluR1) signaling in CNS neurons (Tozzi et al., 2003; Bengtson et al., 2004; Faber et al., 2006; Berg et al., 2007). In substantia nigra dopamine neurons TRPC 5 channels are highly expressed, and mGluR1 agonists induce a current that exhibits a double-rectifying current-voltage plot (Tozzi et al., 2003) similar to the effects of the NKB agonist senktide in Kiss1^ARH^ neurons (**Figure 3**). Both mGluR1 and TacR3 are Gq-coupled to phospholipase C (PLC) activation which leads to hydrolysis of phosphatidylinositol 4,5-bisphosphate (PIP_2_) to diacylglycerol (DAG) and inositol 1,4,5 triphosphate (IP_3_). TRPC channels are minimally Ca^2+^ selective, but can associate with Orai calcium channels to form calcium release-activated calcium channels (Birnbaumer, 2009). A unique feature of TRPC 5 (and TRPC 4) channels is that they are potentiated by lanthanum (La^3+^) (Clapham et al., 2005), which we have exploited to characterize TPRC 5 signaling in POMC neurons (Qiu et al., 2010; Qiu et al., 2014).

Both leptin and insulin excite/depolarize Kiss1^ARH^ and proopiomelanocortin (POMC) neurons through activation of TRPC 5 channels (Qiu et al., 2010; Qiu et al., 2011; Qiu et al., 2014; Kelly et al., 2018). More recently, we documented a critical role of STIM1 in the insulin signaling cascade in POMC neurons (Qiu et al., 2018b). *Stim1* mRNA is highly expressed in POMC (Qiu et al., 2018b) and Kiss1^ARH^ neurons (***Figure 1***), and E2 downregulates *Stim1* mRNA expression in microdissected arcuate nuclei that encompasses these two populations of neurons. Downregulation of *Stim1* is critical for maintaining insulin excitability in POMC neurons with diet-induced obesity (Qiu et al., 2018b). In ovariectomized females that are relatively refractory to insulin excitation, pharmacological blockade of the SOCE complex quickly increases the insulin-mediated excitation of POMC neurons (*i*.*e*., activation of the TRPC 5 mediated inward current), which supports the concept that TRPC 5 channels play a role both in SOCE and receptor operated calcium entry (Birnbaumer, 2009; Salido et al., 2011). Therefore, selective deletion of *Stim1* in Kiss1 neurons should ensure that TRPC 5 channels function as receptor-operated channels to couple TacR3’s and transmit the excitatory effects of NKB to induce synchronous firing of Kiss1^ARH^ neurons as demonstrated in the present findings.

Downregulating STIM1 inhibits SOCE, attenuates Ca^2+^ influx into the ER and elevates intracellular Ca^2+^ levels, which could also contribute to activation of TRPC5 channels in Kiss1^ARH^ neurons (Blair et al., 2009). Indeed, we have found that Ca^2+^ greatly potentiates the leptin-induced TRPC 5 current in POMC neurons (Qiu et al., 2010). In cortical neurons and heterologous cells expressing *Cav1*.*2* (L-type calcium) channels and *Stim1*, inhibition of STIM1 augments Ca^2+^ influx through L-type calcium channels (Park et al., 2010; Wang et al., 2010). Calcium sensing by STIM1 is also involved in the control of L-type Ca^2+^ channel activity in the hippocampal pyramidal neurons such that glutamate-mediated depolarization activates L-type calcium channels, and releases Ca^2+^ from ER stores that activates STIM1 and drives aggregation of the L-type calcium channels to inhibit their activity (Dittmer et al., 2017). Quite possibly in Kiss1^ARH^ neurons and Purkinje cells (Ryu et al., 2017) deletion of Stim1 allows the “dis-aggregation” of TPRC 5 channels, which is reflected in the significant decrease in R_in_ in both cell types with deletion. Furthermore, knocking down STIM1 in cardiomyocyte-derived (HL-1) cells increases the peak amplitude and current density of T-type calcium channels and shifts the activation curve toward more negative membrane potentials (Nguyen et al., 2013). Biotinylation assays reveal that knocking down *Stim1* increases T-type calcium channel surface expression, and co-immunoprecipitation assays suggest that STIM1 directly regulates T-type channel activity (Nguyen et al., 2013). Thus, STIM1 appears to be a negative regulator of voltage-gated calcium channel activity. On the other hand, estradiol treatment in ovariectomized females upregulates *Cav3*.*1* channel expression by 3-fold and whole cell currents by 10-fold in Kiss1^ARH^ neurons, which greatly enhances the excitability and contributes to the synchronous firing of Kiss1^ARH^ neurons (Qiu et al., 2018a). The T-type calcium channel Cav3.1 underlies burst firing in rostral hypothalamic kisspeptin neurons (Zhang et al., 2013b) and facilitates TRPC 4 channel activation in GnRH neurons (Zhang et al., 2008; Zhang et al., 2013a). Cav3.1 channels may also facilitate TRPC5 channel opening in Kiss1^ARH^ neurons (***Figure 8***), but this remains to be determined.

**Figure 8.**
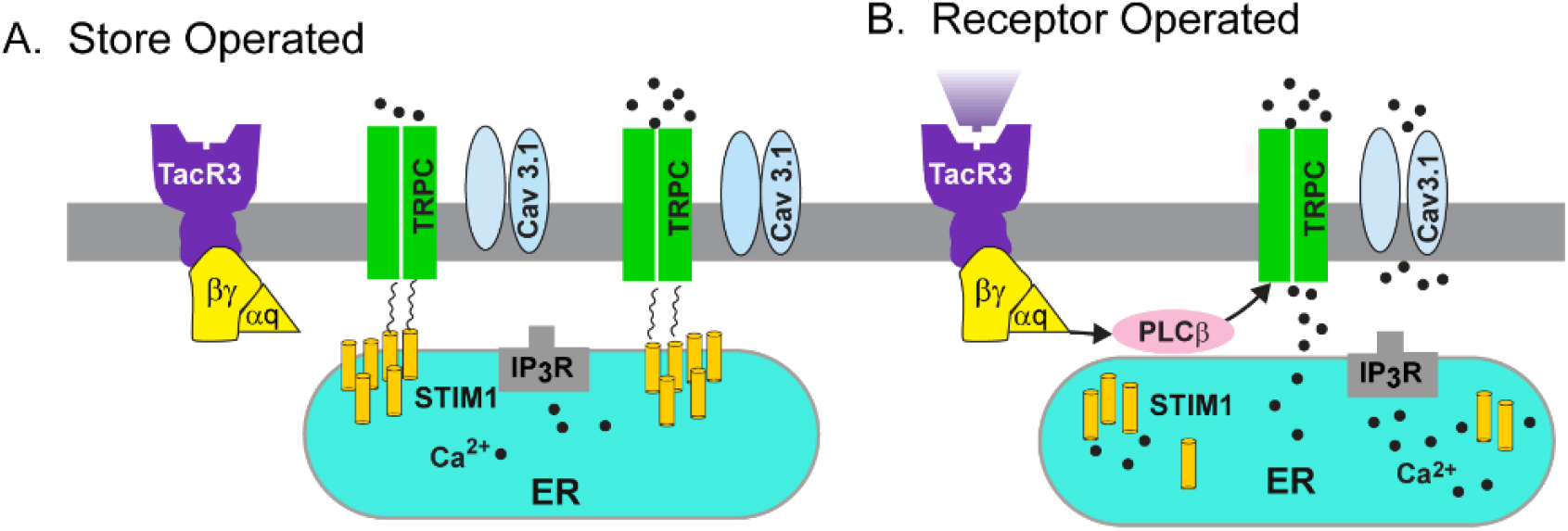
A cellular model of STIM1 affecting NKB activation of TRPC5 channels in Kiss1^ARH^ neurons. **A**, Store-operated calcium entry (SOCE) is a conserved mechanism by which the depletion of the endoplasmic reticulum (ER) is conveyed to calcium-permeable channels at the plasma membrane (PM), triggering calcium influx from the extracellular space and into the cell cytosol. A physiological mechanism responsible for the activation of SOCE results from the stimulation of G-protein coupled receptors associated with the inositol-triphosphate (IP3) and phospholipase C cascade, resulting in the release of calcium from ER, via the IP3 receptor (IP3R). Under physiological stress and in the absence of E_2_, stromal interaction molecule 1 (STIM1) interacts with TPRC5 channels thereby engaging these Ca^2+^ channels as store-operated channels, which are activated with endoplasmic reticulum (ER) depletion of Ca^2+^. **B**, however, under physiological conditions in reproductively active females, in which E_2_ down-regulates the expression of *STIM1*, TRPC 5 channels are converted to receptor-operated channels in Kiss1^ARH^ neurons. Neurokinin B (NKB) binds to its receptor (TacR3) to activate Gαq – PLCβ signaling cascade to facilitate TPRC 5 channel opening, generating a robust inward Na^+^/Ca^2+^ current to depolarize Kiss1^ARH^ neurons, activating T-type calcium (Cav3.1) channels to greatly increase Kiss1^ARH^ neuronal excitability.

Presumably with conditional knockout, *Stim1* was deleted in all cells expressing kisspeptin, which includes arcuate, anteroventral periventricular preoptic (AVPV) and amygdala kisspeptin neurons, and non-neural kisspeptin cells in the gonads, pancreas and liver (Dudek et al., 2019). Currently, we found that the deletion of *Stim1* in kisspeptin neurons had a minor effect on the estrous cycle. *Stim1*^*kko*^ mice exhibited more estrous-type vaginal cytology, which may be indicative of higher levels of circulating estrogens due to increased synchronous firing of kisspeptin neurons and excitatory drive to GnRH neurons (Qiu et al., 2016; Clarkson et al., 2017). It is important to note that synchronous firing of “pulse generator” Kiss1^ARH^ neurons is a failsafe system for maintaining gonadotropin pulses and folliculogenesis in female rodents (Nagae et al., 2021).

Because of the well-documented anorexigenic actions of E2 on POMC and Agouti-related peptide (AgRP) neurons controlling energy homeostasis (Qiu et al., 2006; Roepke et al., 2010; Clegg, 2012; Kelly and Rønnekleiv, 2012; Smith et al., 2013), we ovariectomized the females before feeding them a high fat diet. After 7 weeks on a high fat diet, *Stim1*^*kko*^ females gained modestly less body weight but showed significantly less body fat and lean mass than ovariectomized Kiss1^Cre^ females on a high fat diet. Most importantly, *Stim1*^*kko*^ females exhibited improved glucose tolerance. Kiss1^ARH^ neurons probably mediate these protective effects via their input onto POMC and AgRP neurons. Besides the peptides Kiss1^ARH^ neurons also co-express the vesicular glutamate transporter 2 (vGluT2) (Cravo et al., 2011), and we have documented that optogenetic stimulation of Kiss1^ARH^ neurons expressing channelrhodopsin releases glutamate, which is dependent on the estrogenic state of females (Qiu et al., 2018a). Although the mRNA expression of *Kiss1, Tac2 and Pdyn* mRNA in Kiss1^ARH^ neurons are all down-regulated by E2 (Navarro et al., 2009; Lehman et al., 2010), *Vglut2* mRNA expression is upregulated together with increased probability of glutamate release in E2 treated, ovariectoimzed females (Qiu et al., 2018a). Low frequency (1-2 Hz) optogenetic stimulation of Kiss1^ARH^ neurons evokes fast ionotropic glutamatergic EPSCs in POMC and AgRP neurons, but high frequency (20 Hz) optogenetic stimulation releases enough glutamate to induce a slow excitatory response in POMC neurons but a slow inhibitory response in AgRP neurons (Nestor et al., 2016; Qiu et al., 2016; Qiu et al., 2018a). Indeed, the group I mGluR agonist DHPG depolarizes POMC neurons, while group II/III mGluR agonists (DCG-IV; AMN082) hyperpolarize AgRP neurons (Qiu et al., 2018a). Group I mGluRs (mGluR1 and mGluR5) are G_q_/G_11_-coupled, while group II/III mGluRs (mGluR2 and mGluR7) are G_i_/G_o_-coupled (Niswender and Conn, 2010). Hence, the output of Kiss1^ARH^ neurons excites the anorexigenic POMC neurons and inhibits the orexigenic AgRP neurons. Therefore, Kiss1^ARH^ neurons appear to be an integral part of an anorexigenic circuit in the hypothalamus (Qiu et al., 2018a; Rønnekleiv et al., 2019; Navarro, 2020).

Presently, there is compelling evidence that Kiss1^ARH^ neurons are a critical “command” neuron for coordinating energy states with reproductive functions (see (Rønnekleiv et al., 2019; Navarro, 2020) for review). We have now documented that conditional knockout of *Stim1* in Kiss1^ARH^ neurons, which augments the NKB-mediated depolarization of these neurons via TRPC 5 channels, helps protect ovariectomized, female mice from diet-induced obesity and glucose intolerance. In addition, in preliminary experiments we have found that insulin treatment *in vitro* increases the synchronous firing (GCaMP6 activity) of Kiss1^ARH^ neurons, which further emphasizes its role as a “command” neuron. Clearly, Kiss1^ARH^ neurons are part of a hypothalamic circuit for coordinating reproduction with energy balance, but additional experiments are needed to elucidate the cellular mechanisms by which steroid and metabolic hormonal signaling synergize to govern their activity.

## Acknowledgements

This research was funded by National Institute of Health (NIH) grants: R01-NS043330 (OKR), R01-NS038809 (MJK) and R01-DK068098 (OKR and MJK). Confocal microscopy was supported by a P30 NS061800 (PI, S Aicher) grant. We thank Mr. Daniel Johnson for his technical support.

---

**Figure 2-video supplement 1. Neurokinin B receptor agonist senktide induces [Ca**^**2+**^]_**i**_ **increase in Kiss1**^**ARH**^ **neurons expressing GCaMP6s**. Imaging of transient Ca^2+^ changes in an arcuate slice using spinning disk confocal microscopy. Fluorescence intensity was measured over 20 minutes, before and after application of senktide (1 µM). The period represented is 20 minutes.

**Figure 3-video supplement 1. High frequency photo-stimulation induces a slow excitatory postsynaptic potential (slow EPSP)**. Slow EPSP was induced by a 10-s 20 Hz photostimulation (light intensity 0.9 mW and pulse duration, 10 ms) in a ChR2-expressing Kiss1^ARH^ neuron in a slice from a *Kiss1*^*Cre*^*::Ai32* mouse. The period represented is 1 minute, 34 seconds.

## Notes

### Competing Interest Statement

The authors have declared no competing interest.

